# Contrast-Enhanced, Molecular Imaging of Vascular Inflammation in the Mouse Model by Simultaneous PET/MRI

**DOI:** 10.1101/2019.12.16.878652

**Authors:** Siyi Du, Thomas S.C. Ng, Adrian House, Tang Tang, Lin Zheng, Chuqiao Tu, Janice Peake, Imelda E. Espiritu, Kwan-Liu Ma, Kent Pinkerton, Russell E. Jacobs, Angelique Y. Louie

**Affiliations:** Chemistry Graduate Group, University of California, Davis, United States of America; Division of Biology & Biological Engineering, California Institute of Technology, United States of America; Department of Biomedical Engineering, University of California, Davis, United States of America; Department of Computer Science, University of California, Davis, United States of America; Department of Anatomy, Physiology and Cell Biology School of Veterinary Medicine, University of California, Davis, United States of America

**Keywords:** Multimodality imaging, MRI/PET, dual-mode imaging, nanoparticle, atherosclerosis, vulnerable plaque, cardiovascular imaging, vessel wall, VCAM

## Abstract

Despite advances in diagnosis and management, cardiovascular diseases (CVDs) remain the leading cause of death in the US. Atherosclerosis is the most common form of CVD and the vulnerability of atherosclerotic plaques to rupture is a primary determinant for risk of catastrophic ischemic events. Current imaging of atherosclerotic disease focuses on assessing plaque size and the degree of luminal stenosis, which are not good predictors of plaque stability. Functional methods to identify biomarkers of inflammation in plaques could facilitate assessment of plaque instability to allow early intervention. In this study, we validate the use of a purpose-built, magnetic resonance imaging (MRI)-compatible positron emission tomography (PET) insert for multimodal, molecular imaging of vulnerable plaques in mice. We illustrate the application of PET to screen for inflamed regions to guide the application of MRI. Molecular MRI visualizes regions of vascular inflammation and is coupled with anatomical MRI to generate detailed maps of the inflammatory marker within the context of an individual vessel. As a testbed for this imaging methodology, we developed a multimodal, iron oxide nanoparticle (NP) targeting vascular cell adhesion molecule-1 (VCAM-1) for simultaneous PET/MRI of vascular inflammation performed on a mouse carotid ligation model. *In vitro* cell studies confirmed that the NPs are not cytotoxic to liver cells. *In vivo* simultaneous PET/MRI imaging identified regions of inflammation. Three-dimensional rendering of the MRI data facilitated high-resolution visualization of patterns of inflammation along the injured vessel. Histology validated the co-localization of the NPs with VCAM-1 expression at sites of induced inflammation. The results of this work validate the utility of the simultaneous PET/MR insert as a research tool for small animals and lays groundwork to further advance the potential clinical utility of integrated imaging systems.

## 1. Introduction

Cardiovascular disease is the leading cause of death for both males and females in the United States.[1] In particular, atherosclerosis is responsible for catastrophic manifestations of heart disease such as myocardial infarction and stroke.[2] Recent understanding of the pathophysiology of plaque formation has identified chronic inflammation as a hallmark of plaque development; localized inflammatory response can lead to the development of “vulnerable” plaques that are prone to rupture and cause downstream vascular occlusion.[3] Imaging can play a role in identifying patients with vascular lesions susceptible to acute cardiovascular events, who may be amenable to treatment with anti-inflammatories or interventional procedures. Recent literature has shown that plaque lesion composition, particularly the presence of inflammatory markers and immune cells, as opposed to the degree of vessel stenosis, is a better predictor of patient mortality and morbidity; and assessment of plaque inflammation is an excellent target for noninvasive imaging.[4] However, current clinical imaging techniques seldom provide specific information about inflammation.

Current clinical imaging techniques such as coronary angiography, vascular ultrasound and computed tomography focus on identifying stenotic disease and can miss vulnerable plaques that do not cause significant structural stenosis.[5] Anatomical features identified by coronary computed tomography (CT) and magnetic resonance imaging (MRI) that could be used to classify plaques, have not been fully validated for prediction of vulnerability.[6] Targeted molecular imaging has potential for greater predictive value. For example, imaging of plaque inflammation has been actively pursued using ^18^F-labeled fluorodeoxyglucose (FDG) positron emission tomography (PET) and has shown promise for identifying inflammation, generally in large vessels such as the carotid and aorta.[7–10] However, it is challenging to accurately assess inflammatory burden in small vessels with the limited resolution of PET.[11] Moreover, FDG is a non-specific marker of inflammation that measures only glucose uptake, which presents challenges for coronary artery imaging against the high metabolic background of the myocardium.[12] Enthusiasm for the clinical use of ^18^FDG to identify plaques has diminished over recent years with new tracers such as ^18^NaF receiving greater attention; but specific imaging of plaque inflammation is still not available.[13] New, targeted imaging strategies that can identify inflamed plaques in smaller vessels, such as the coronary arteries, with higher specificity are needed.

We have previously shown that multimodal agents combining positron emission tomography (PET) and magnetic resonance imaging (MRI) can detect macrophage density in plaques, using the quantitative ability of PET for high sensitivity mapping of the cells, and MRI for high spatial resolution molecular imaging and soft tissue mapping.[14] Our previous work utilized separate scanners to visualize the synergistic PET and MRI information. This required laborious spatial co-registration (prone to misalignment since they are acquired sequentially), more involved handling of subjects with longer sedation, and time delay due to the need for transport between PET and MR scanners. The latter factor also prevents temporal co-registration of the PET and MRI signals. Not only does this hinder throughput of preclinical research pursuits, but also may limit the translatability of such agents to the clinic. The use of hybrid modality instruments for simultaneous signal acquisition is, thus, of increasing interest in the imaging field and clinical hybrid instruments are now available; but the ideal cardiovascular applications for these hybrid systems are still under investigation.[15]

In this proof of concept study, we demonstrate the utility of simultaneous PET and MRI to facilitate PET-guided MRI mapping of localized inflammation in small vessels using an integrated small animal simultaneous PET/MRI imaging system developed at UC Davis and a dual-mode contrast agent targeted to Vascular Cell Adhesion Molecule 1 (VCAM-1).[16, 17] Vascular Cell Adhesion Molecule VCAM-1 has received attention for imaging of atherosclerotic plaques due to its overexpression in the pathogenesis of vulnerable plaques. It has been studied for single modality imaging by MRI[18], SPECT[19, 20], and ultrasound[21]. Each of these modalities holds inherent limitations for targeted vascular imaging. MRI and ultrasound can provide excellent local, spatial information at the lesion site, but are ill-suited to whole-body screening. PET is superior for screening, but lacks the spatial resolution to identify individual vessels. While computed tomography (CT), can provide structural information, but does not provide physiological information at the lesion site. MRI/PET has the potential to overcome these difficulties by combining the screening power of PET with the resolution power of MRI, enabling concurrent assessment of cardiovascular structural and physiological abnormalities.[22] We evaluated the ability of the integrated PET/MRI instrument to visualize sites of localized inflammation induced by carotid injury in a mouse model; the mouse carotid is of similar size to human coronary arteries. PET provided an overview of inflamed regions, which was used to focus MRI interrogation and obtain higher resolution, 3-dimensional details of VCAM-1 expression in single vessels.

## 2. Results and Discussion

Nanoparticles (NP) were successfully synthesized (outlined in Figure 1 and described in Methods).

**Figure 1.**
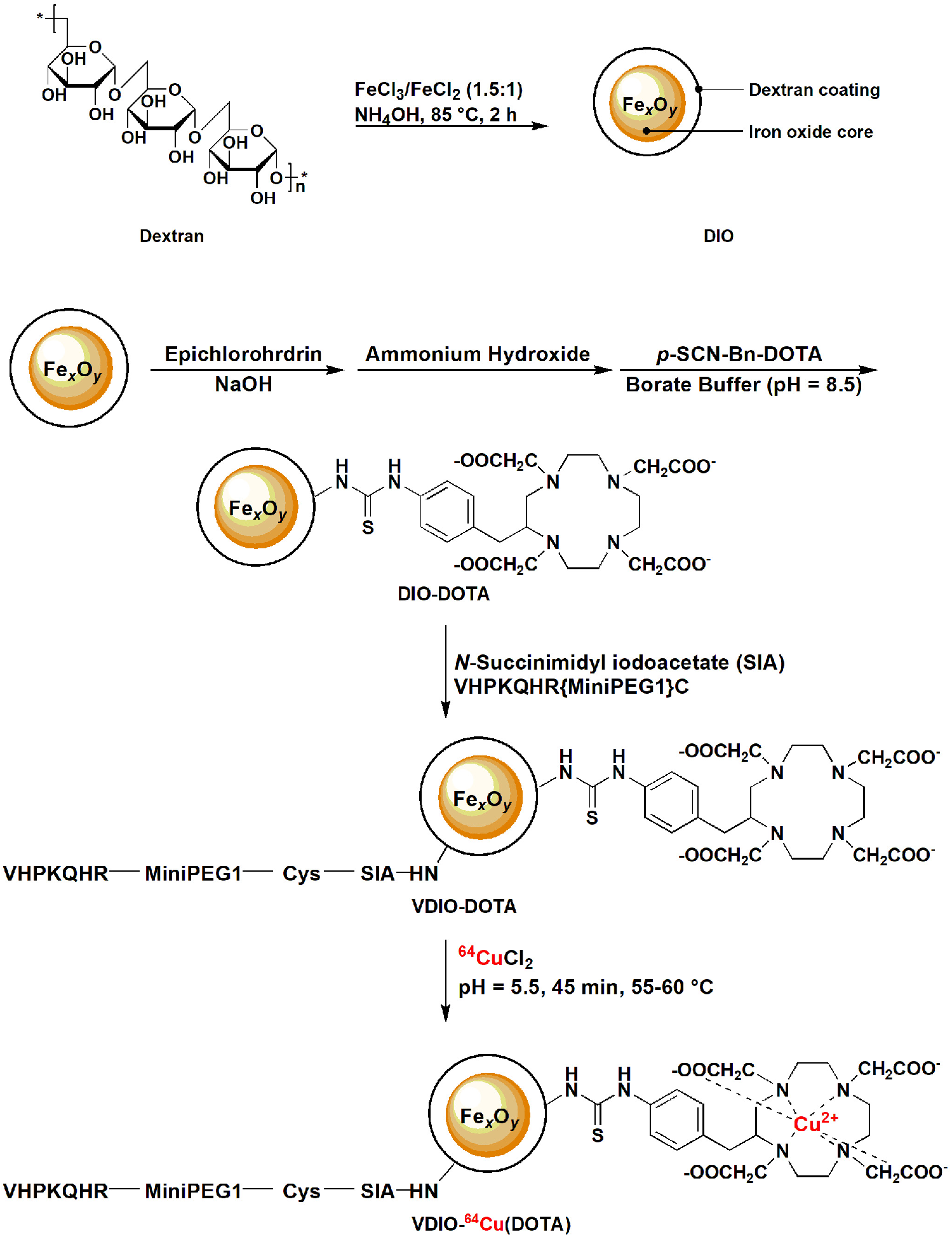
Synthesis of VDIO-DOTA nanoparticles. (a) Synthesis of dextran coated iron oxide nanoparticle (b) Conjugation of DOTA and VCAM-1 targeting peptides to nanoparticles.

### 2.1 Characterization of VDIO-DOTA and DIO (control)

DIO (dextran-coated iron oxide) was synthesized as a matched control to the prepared VDIO-DOTA (DOTA=1,4,7,10-Tetraazacyclododecane-1,4,7,10-tetraacetic acid; VDIO=VCAM-conjugated DIO;). The physical properties of VDIO-DOTA and its precursor DIO are summarized in Figure 2a. Following the syntheses described, the DIO and VDIO-DOTA iron oxide core sizes were measured to be 7.3 ± 2.9 nm by averaging 500 particle measurements from TEM images as shown in representative images in Figure 2b and 2c. Inset plots show that the hydrodynamic diameters of DIO and VDIO-DOTA were found to be 39.7 ± 15.0 nm and 48.1 ± 19.5nm, respectively, using dynamic light scattering (DLS). The increased hydrodynamic diameter could be explained by the addition of peptide polymers on the VDIO surface holding the dextran polymers apart via steric hindrance. However, the core sizes remained the same.

**Figure 2.**
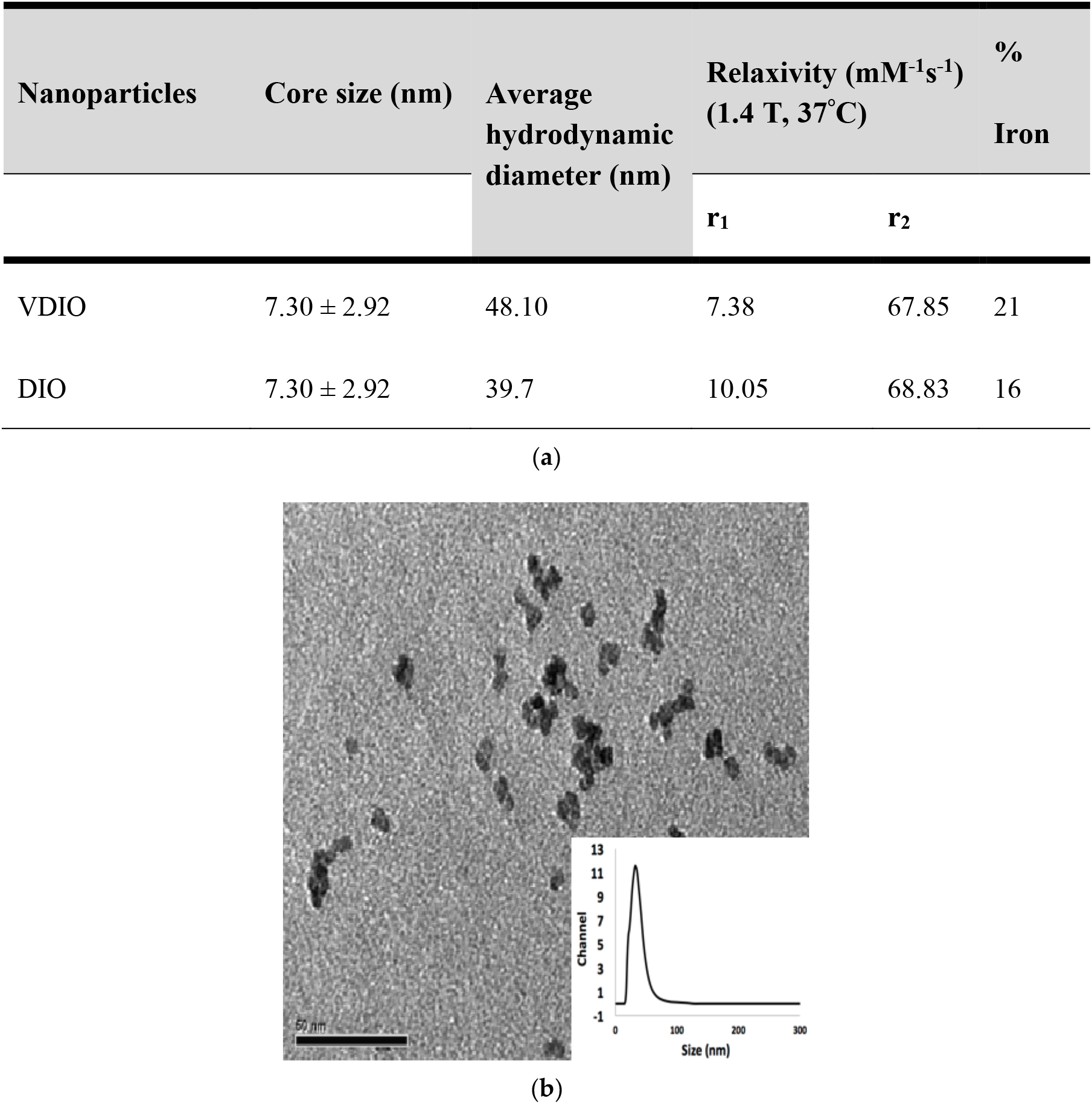
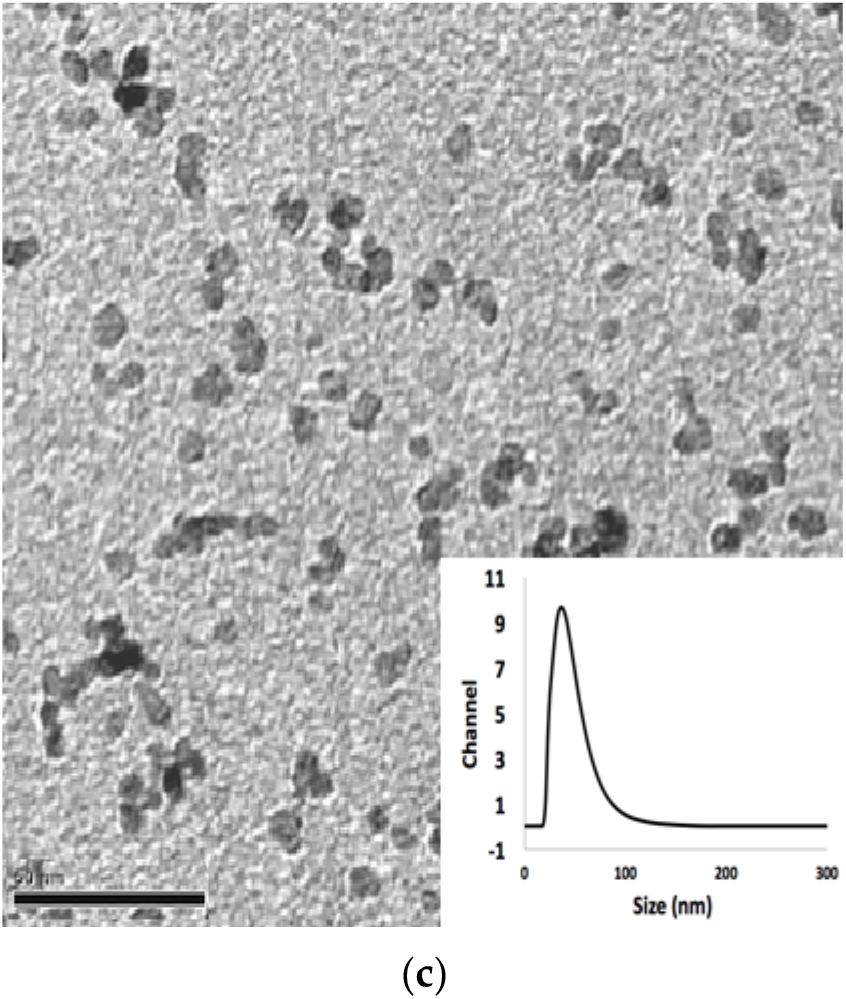
(a) Summary of nanoparticle properties. TEM images of (b) DIO (core size: 7.30 ± 2.92nm) (C) VDIO-DOTA (core size: 7.30 ± 2.92nm). Scale bars = 50 nm, insets are DLS data.

**Figure 3.**
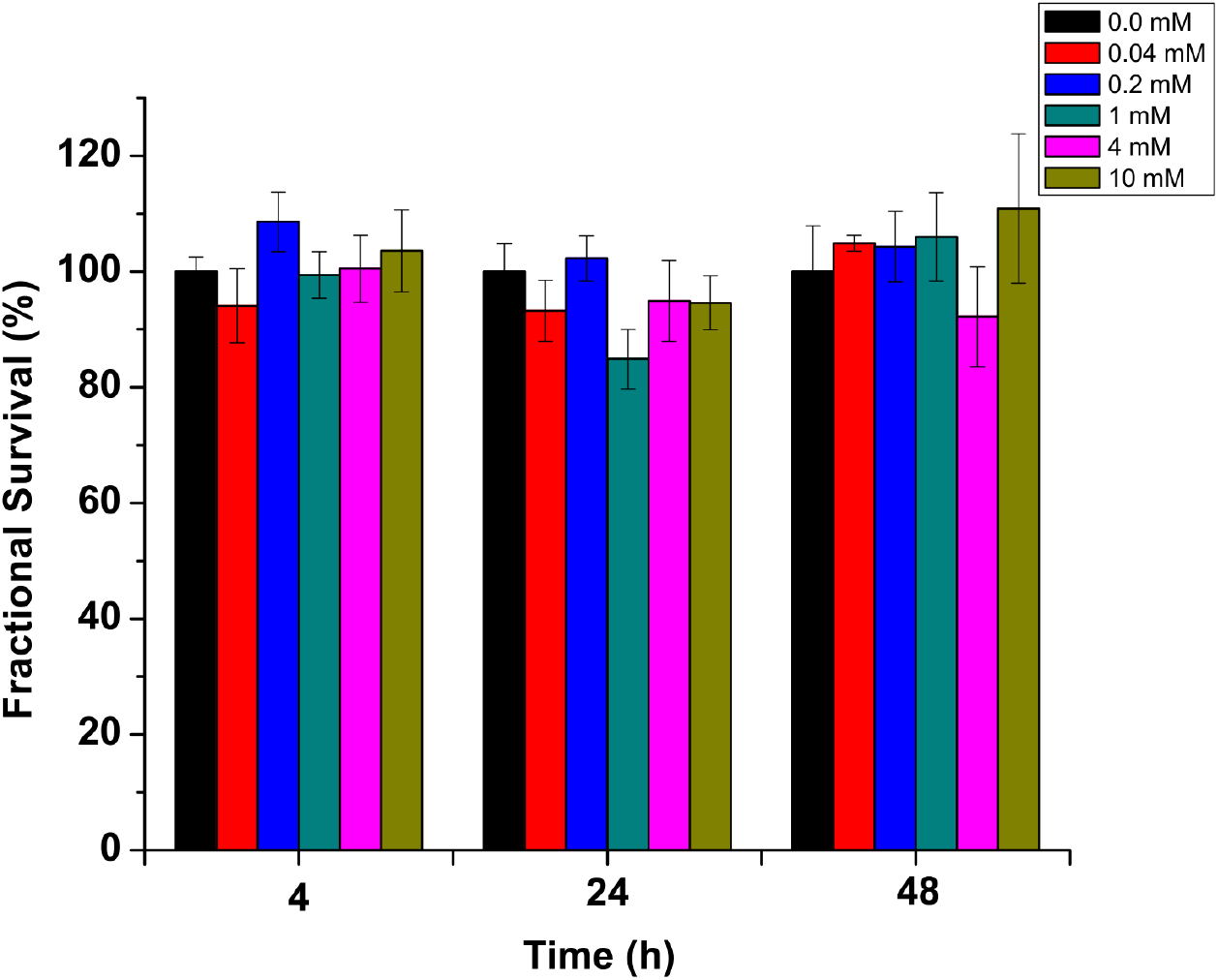
The fractional survival of HepG2 after 4, 24 and 48 hours of incubation with VDIO solutions of varying concentrations. At each time point, fluorescent intensities reflecting survival fractions (y axis) were normalized against the signal from the untreated control group (black bar at each time point).

The iron concentration (mg per unit mass) in DIO and VDIO-DOTA was measured by atomic absorption (AA) to be 0.161 and 0.037 mg Fe/mg of nanoparticles, respectively. The longitudinal (r_1_) relaxivity of VDIO-DOTA at 60 MHz (1.4 T, 37 °C, pH = 7) was 7.4 mM^−1^s^−1^ and transverse relaxivity (r_2_) was measured as 67.9 mM^−1^s^−1^. DIO had r_1_ and r_2_ values of 10.1 mM^−1^s^−1^ and 68.8 mM^−1^s^−1^, respectively. The r_2_ to r_1_ ratio for both reflects their suitability as T_2_-weighted MRI contrast agents.[17] VDIO-DOTA relaxivity and size remained stable after 1 year of dry storage at room temperature.

### 2.2 Contrast Agent is Not Cytotoxic to Liver Cells in vitro

Given that NPs are expected to clear through the liver, liver cells may be exposed to the highest off-target concentration of injected contrast agent. Thus, C_12_ – Resazurin viability assays were performed on HepG2 liver cells to evaluate any toxicity exhibited by VDIO-DOTA.[23] In this assay resazurin is reduced to resorufin in proliferating cells.[23] Cell survival was evaluated after incubation for 4, 24, and 48 hours with different concentrations of VDIO-DOTA, ranging from 0.04 mM iron (red), 0.2 mM iron (blue), 1 mM iron (teal), 4 mM iron (pink) to 10 mM iron (khaki). With 90% confidence interval by t-test there were no differences between the untreated control (0.0 mM iron, black) and cells treated at all concentrations of VDIO-DOTA tested from 0.04 mM iron up to 10 mM iron. These results support that the nanoparticles are nontoxic to liver cells at relatively high concentrations.

### 2.3 VDIO-DOTA can to detect inflamed vessels in vivo

MRI only studies were performed to determine appropriate dosing prior to PET/MRI studies. Representative MRI scans showing VDIO-DOTA uptake at the three injection concentrations are shown in Figure 4a (n = 4 mice per concentration). The leftmost column shows cross sections through the entire animal for, from top to bottom 6mg/ml, 30mg/ml and 60 mg/ml injection of nanoparticles. These are labeled with colored circles to indicate the injured vessel (yellow circle), uninjured contralateral vessel (red circle) and spinal cord (green circle). The co-registration standards are also visible to left and right of the animal. The regions outlined by the yellow boxes are presented in zoomed views on the matrix of images on the right, which show, from left to right, images from this region taken pre-injection, and images 4 and 24 hours after injection of contrast. For all concentrations, the inflamed vessel (yellow arrows) showed expected increased dephasing (seen as signal dropout) as the concentration of VDIO-DOTA injected is increased resulting in greater accumulation at the ligated site compared to the nonligated contralateral vessel (red arrows). This darkening in the inflamed arteries persists through the 24h timepoint for all concentrations, showing sustained retention of the particles at the site of inflammation. Robust signal for 6ml/ml suggested that even lower injection concentrations could be employed. The CR was calculated as a function of time after injection and dosage and shown in the accompanying graph in Figure 4b. Although the 60 mg Fe/kg dose showed the greatest signal decrease on the images, maximal CR was achieved with the 30 mg Fe/kg dosage at both 4h and 24h after injection of contrast. This highlights the need to achieve a balance between local accumulation of the NP, retention over time, and its systemic circulation for optimal visualization of the region of interest. Using these results as a guide, 30mg Fe/kg was used for the remainder of the studies.

**Figure 4.**
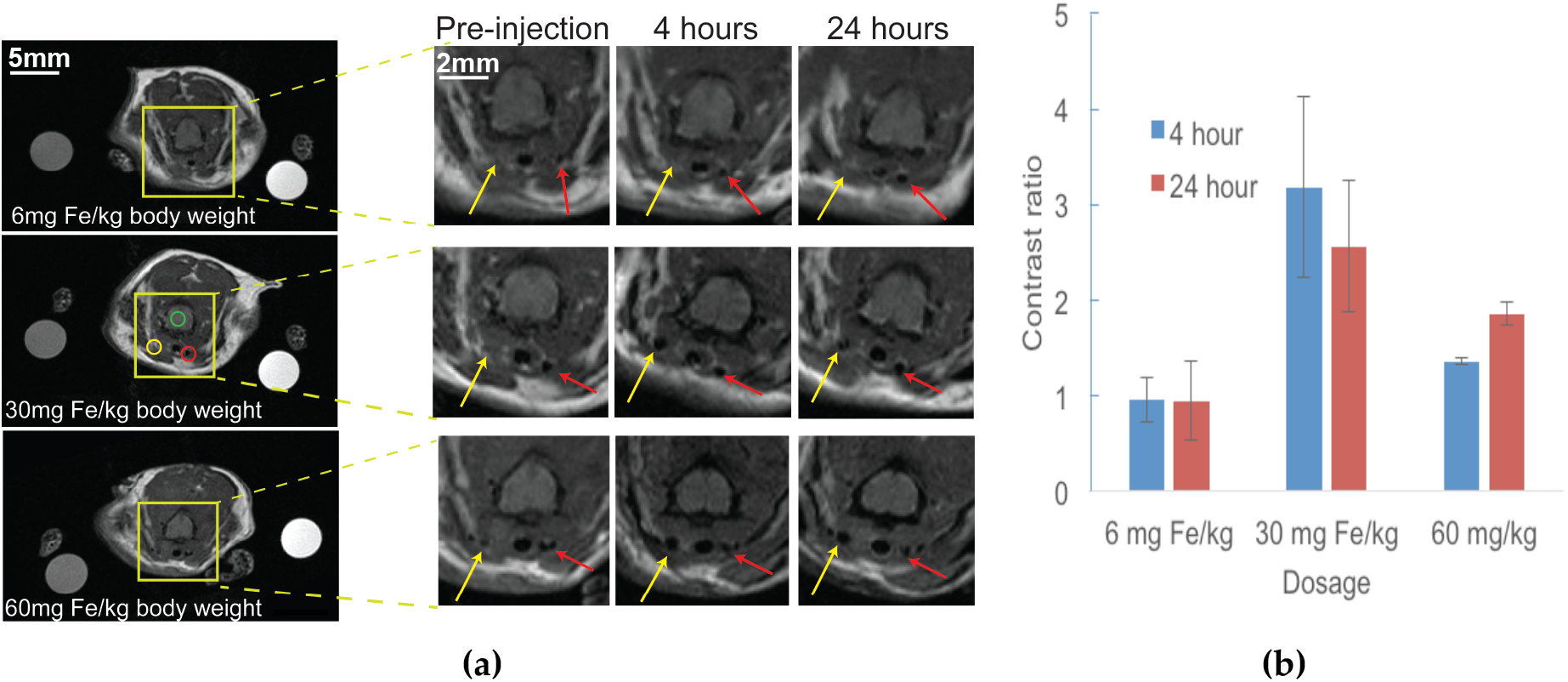
(a) MRI-only study showing VDIO-DOTA uptake at different injection concentrations over 24 hours. The yellow arrows point to the inflamed vessel, while the red arrows indicate the non-inflamed control vessel. There was signal decrease at the site of injury at all dosages; but the decrease was most prominent at the 30 and 60 mg Fe/kg. The regions of interest for Contrast Ratio (CR) calculation are shown by the circles. (Scale bar = 5 mm). (b) Contrast ratio as function of time and dose for MRI-only studies. Compared to baseline, the 6 mg Fe/kg dose level did not demonstrate significant local accumulation of NP at the site of inflammation. Both the 30 mg Fe/kg and 60 mg Fe/kg showed increased accumulation of NP at the site of inflammation; the 30 mg Fe/kg dose demonstrated the highest CR, likely due to the reduction of signal differentiation between the localized accumulation and systemic distribution of NP at the 60 mg Fe/kg dosage. Thus the 30 mg Fe/kg dose was used for subsequent studies. (Error bars denote the SEM).

Following these MRI only studies, we evaluated VDIO-^64^Cu-DOTA nanoparticle accumulation *in vivo* by hybrid PET/MRI. Figure 5 shows image slices by MRI (top row), by PET (middle row) and the overlay (bottom row) from the same cross section in a representative mouse (n = 4 mice total). Prior to injection of VDIO-^64^Cu-DOTA, no MR or PET signal beyond background levels was seen within the animal. At 4 h post injection, both carotids demonstrated PET signal, which was higher in the injured vessel, suggestive of radiolabeled NP accumulation at the site of injury along with continued systemic circulation of unbound NP. Another focus of PET uptake within the field of view appear to correspond with a vein, supporting that systemic circulation of the NP remains at this time point, but which could represent another site of inflammation in the animal. In the slice shown, at 4 h there is a signal increase in the ligated vessel (yellow arrow) at a ratio of 1.19 compared to the uninjured contralateral vessel; at the 24 h time point the ratio of the ligated site compared to the contralateral artery was 1.07; partial volume effects limits the accuracy of these measurement. We also calculated the MRI contrast ratio (CR; see equation in methods 4.6.3) from the concurrent MRI dataset. Note that CR also is a comparison against the contralateral control, thus taking into account contributions from signal in the blood. For this particular animal, the MRI CR was 1.41 at the 4 h time point and 0.62 by 24 h, indicating elevated NP accumulation at the ligated site at the 4 h timepoint with decreased MRI signal by 24 h, commensurate with the PET results. This slice was selected to illustrate the point that these trends are for this slice, in this particular animal and based on this slice only one may conclude there was no inflammation in this vessel. For a more accurate view of inflammation the entire affected volume should be considered, as well as the full set of experimental subjects. As shown later, mapping inflammation throughout the vessel provides a clearer picture of inflammatory burden. The MRI CRs for the entire PET/MRI cohort were 1.65 ± 0.26 at 4 h and 1.66 ± 0.39 at 24 hours respectively. In general, this is in agreement with the data for the MRI only cohort, which also demonstrated an increased MRI CR over the 24 h period (Figure 5b). Note that these results support that ligation does not prevent contrast agent access to the injured carotid—if this had been the case there would be less signal drop out due to reduced perfusion to the region.

**Figure 5.**
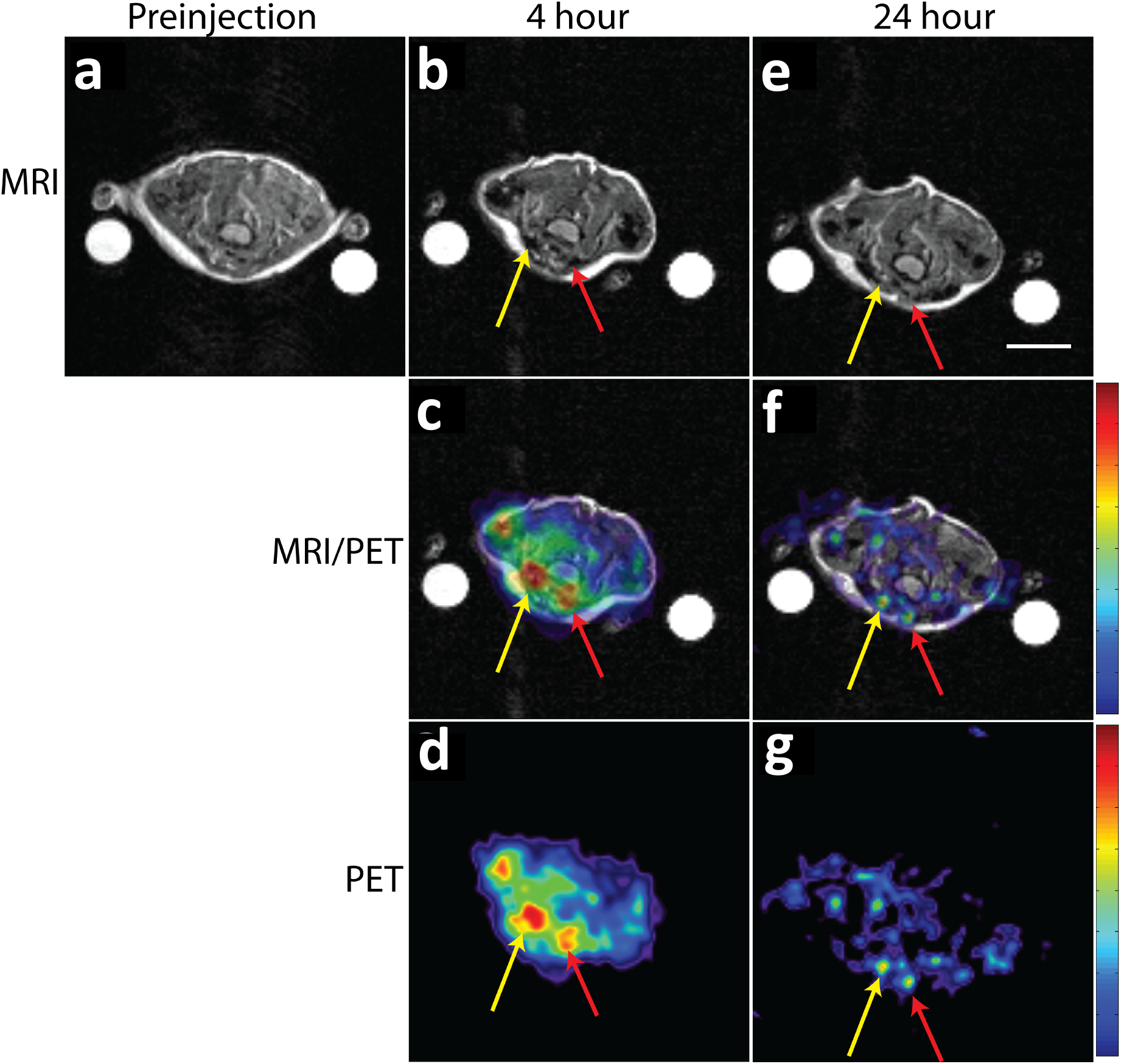
PET/MR imaging of dual-mode VDIO-DOTA accumulation in carotid artery of ApoE^−/−^ mouse with vascular inflammation (atherosclerotic plaques) induced by ligation. MRI-only images preinjection (a), after 4h(b) and 24h(e) injection, PET/MRI co-registered images after 4h(c) and 24h(f) injection and PET-only images after 4h(d) and 24h(g) injection are shown. The image slices were from the same cross section of the mouse and showed spinal cord (grey oval area in the middle), left carotid (after ligation) artery (denoted by yellow arrow) and right carotid artery (denoted by red arrow). The color maps for the PET images were set such that the highest values (as indicated by red) reflected the highest signal within the animal image volume for the particular time point. The white circles are markers for MRI intensity standards. (Scale bar = 5 mm).

The lower resolution of PET provides decreased accuracy in determining inflamed regions due to partial volume effects, and subtle inflammatory lesions may be missed. This is suggested by the fact that although analysis of the MRI data suggested there was accumulation of the NP at the ligated site, this was not reflected by the simultaneously acquired PET data. The ratios of the PET signal at the ligated artery compared to the control artery were 1.05 ± 0.04 at 4 hours and 0.82 ± 0.33 at 24 hours. This highlights the need for high-resolution 3D renderings of the affected volume to better identify regions of high inflammatory activity and assess degree of instability. For improved volume visualization, we performed three-dimensional rendering of the MRI data as shown in Figure 6. MRI rendering is able to define regional accumulations and concentration differences for VDIO-DOTA in the vessel wall, allowing us to better localize VCAM-1 expression to specific regions in the inflamed vessel. Figure 6a and b show rendering of the MRI only data from the 30 mg Fe/kg dose injection for a representative animal, with a slice from the anatomical MRI included for anatomical context; this is the same animal and slice view used for the cross section image shown in Figure 4. The color map (transfer function) for the image was set to define three major color zones representing low (red), medium (blue), and high (pink) NP uptake. Greater susceptibility from T_2_-weighting indicates regions of greater particle accumulation (more red). The volume renderings suggest that VCAM-1 expression, which is targeted by the NPS and indicative of activated endothelial cells and inflammation, is diffuse along the vasculature and that expression patterns vary across the vessel. In this particular animal there is a greater accumulation of contrast agent on one side of the vessel as seen in the 4 h still images in panel a. At 24h (panel b) the trend persists and one can observe accumulation on one side that is consistent with the 4-hour data.

**Figure 6.**
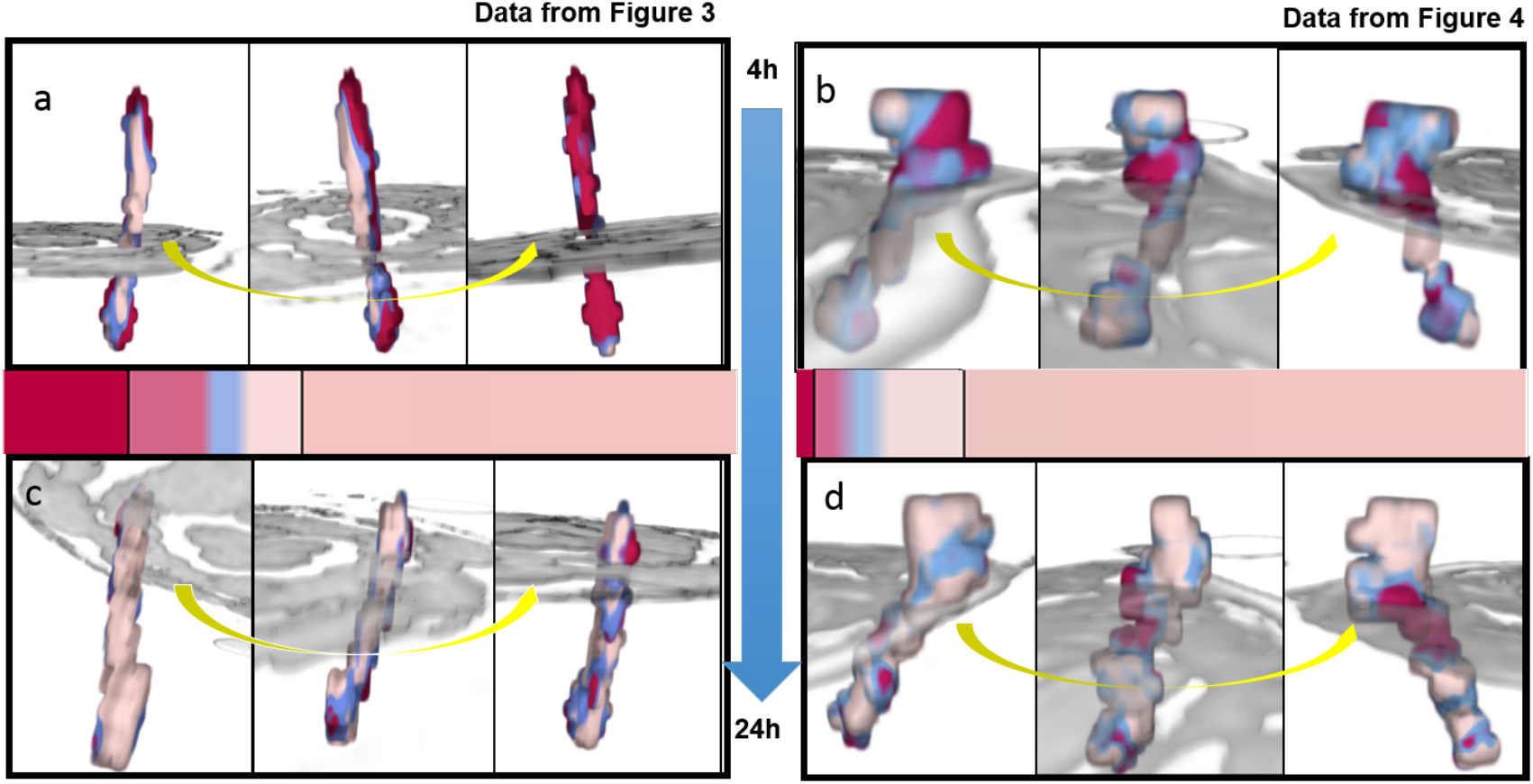
Three-dimensional rendering of MRI data (a – b). Medium dose study, MRI data (from Figure 3). (a) Still frames from video depicting a rotating view of the inflamed vessel 4 h after injection of contrast agent. (b) Similar data for the vessel at 24 h. A similar trend for higher accumulation of contrast agent on one side of the vessel is preserved at both time points. (c – d) PET/MRI study, MRI data (From Figure 4). (c) Still frames from video depicting a rotating view of the inflamed vessel 4 h after injection of contrast agent. (d) Similar data for the vessel at 24 h. A trend for a “hot spot” (red) accumulation of contrast agent at the same location is seen at both time points. Transfer functions used for color assignments are displayed as color bars shown between the 4 and 24h data sets.

Figures 6c and d display rendering of the MRI data for the animal shown in the PET/MRI images of Figure 5. Inflammation is even more heterogeneous in this animal and NP accumulates in patches throughout the vessel. A local “hot spot”, appears in the 4 h data (panel c) and persists but is decreased through the 24 h time point (panel d) commensurate with the PET and MRI CR results. At 24 h most of the hot spot lies below the plane of interrogation that was show in the slice in Figure 4; this may explain the lack of PET observed in the slice shown and also underscores the value of evaluating 3D data sets as volume renderings. The rendering provides insight into the heterogeneity of the NP accumulation, which is not readily apparent from slice by slice visualization of the MRI image set. This data correlates with the histological data, described in the next section, which showed inhomogeneous regional accumulations of the nanoparticles that overlap with regions showing high VCAM-1 expression. The morphology of the local plaque load can have prognostic implications.[24] Thus, this additional information from the high resolution MRI may provide vital information for clinical management.

The addition of a 3D visualization is very useful for interpreting these types of data sets where the disease is diffuse and lesions can span across multiple slices. One of the challenges for 3D rendering is the ability to create images and videos without the need for extensive manual segmentation, i.e. user delineation of pathology. The renderings shown here were created using software created at UC Davis that performs intensity-based segmentation by applying a user-defined color map to all voxels in the data set. This removes some degree of subjectivity from the assignment of pathological features.

### 2.4 Iron co-localizes with VCAM-1 expression

Mice were sacrificed after *in vivo* MRI only or PET/MRI study and organs of interest were collected for further analysis. For the MRI only studies, the tissues from mouse carotids were embedded into paraffin sections for histological study to examine iron accumulation (Figures 7a, c, e) and VCAM-1 expression (Figures 7 b, d, f) in the tissues. Prussian Blue staining was used to detect ferric ion (Fe^3+^) in the tissue. As shown in Figure 7a blue staining demonstrated that iron could be found in the intima in the inflamed carotid artery (arrow). Anti-VCAM-1 antibody staining was performed on an adjacent section (Figure 7b), which confirms the presence of VCAM-1 in the same region, as indicated by dark brown stain of the intima on the left side of the vessel lumen (arrow). The dark blue iron staining co-localized in the regions of darkest brown VCAM-1 staining, supporting that the nanoparticles were co-localized with VCAM-1 expression. There appears to be a thin layer of VCAM-1 staining on the rest of the lumen that does correlate with some iron staining. The faint brown stain over the entire intima represents nonspecific staining that is also found in the control vessel. Non-inflamed contralateral vessel showed no iron (Figure 7c) or dark brown VCAM staining (Figure 7d).

**Figure 7.**
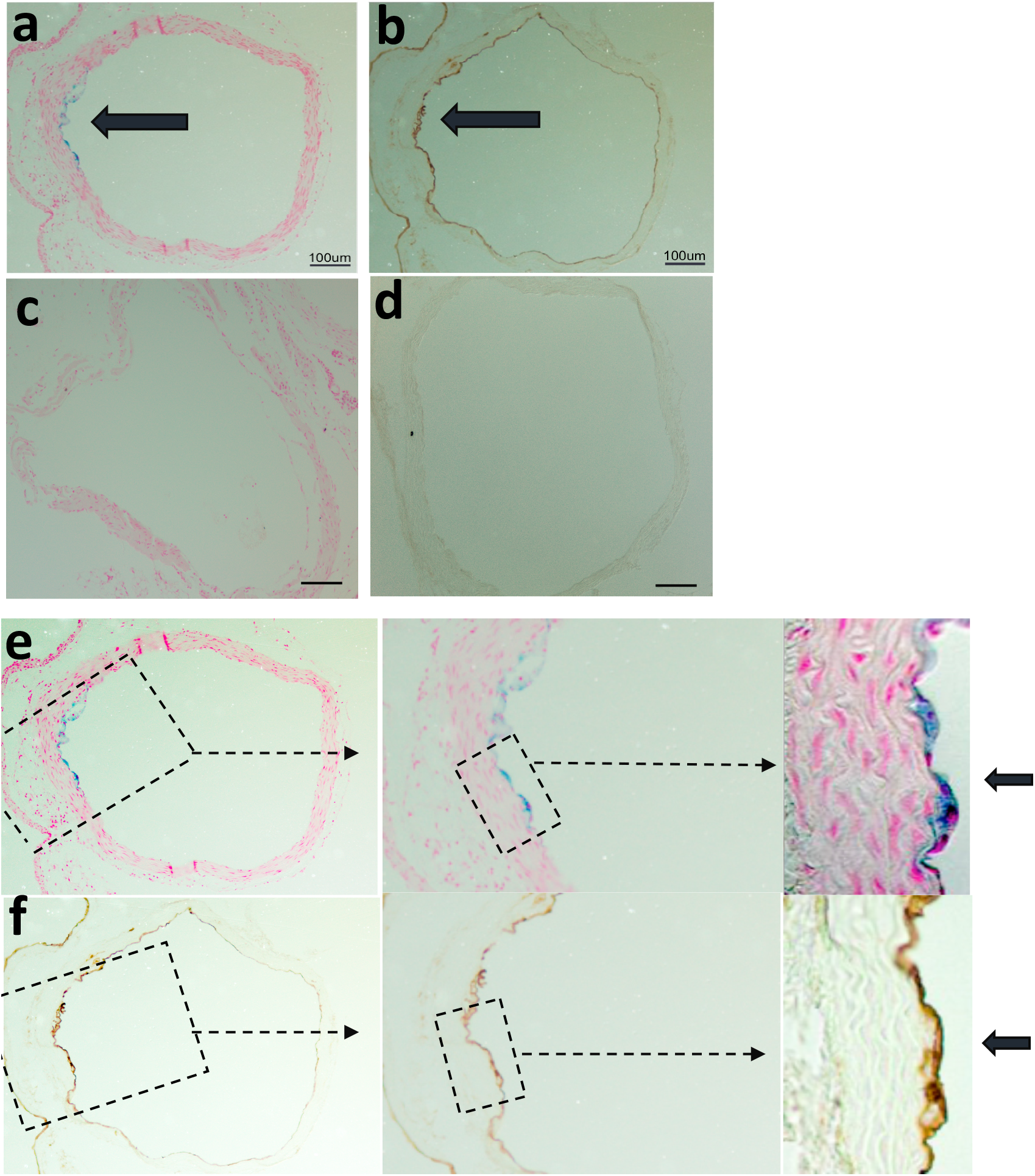
Iron colocalizes with VCAM-1 expression. (a). Prussian Blue staining hows iron on tissue cross section from inflamed mouse carotid artery. (b). Immunohistochemical staining shows VCAM-1 (brown) on a tissue slice adjacent to that shown in A. (c). Prussian Blue staining showed no iron in tissue cross section from normal mouse carotid artery and (d) no VCAM-1 staining on neighboring tissue slice. (e) Higher magnification view of Prussian Blue staining in panel a at 10X, 20X and 40X. (f) Higher magnification views of VCAM-1 staining in panel b. Dashed boxes indicate the area of interest with increasing magnification, moving left to right. Staining appears localized to the monolayer of endothelial cells lining the vessel.

VCAM-1, triggered by inflammatory cytokines such as TNF-α, is expressed by endothelial cells as a prelude to the recruitment of macrophages into plaques. In our previous work to label macrophages in the injury model, animals were placed on fat diet for 2 weeks after ligation; the current studies were performed 8 days after ligation. At this earlier time point, plaque size and macrophage involvement are minimal as can be seen in the higher magnification histological images. Figure 7e shows iron staining at increasing magnifications from left to right and iron staining localized to cells of the intima, which is still generally a monolayer of endothelial cells. Figure 7f shows VCAM-1 staining at increasing magnifications from left to right, which shows VCAM-1 localized to the same monolayer. There was no intimal thickening, or obvious macrophage accumulation observed in the histological slices. The iron stain also appeared confined to a monolayer of “cobblestone-like” endothelial cells and there was no evidence of macrophages, which display surface ruffles and blebs. These results support that the NPs were binding to VCAM-1 expressed on endothelial cells.

## 4. Conclusions

In this study, we evaluated the use of a simultaneous hybrid PET/MRI system and multimodal nanoparticle contrast agent to visualize distribution of an early marker of inflammation. We found robust spatial-temporal co-localization of the PET and MRI signal, and identified regions of inflammation, which were confirmed by histological staining. PET, limited by its lower spatial resolution, could not specifically map inflammation in the vessels. PET also missed elevated contrast that was detected by MRI, likely due to signal averaging effects of diffuse inflammation in the tissues. But the sensitivity of PET allowed detection of inflammation in a larger volume map and served to guide MR imaging. Careful analysis of the MRI data, aided by 3D volume rendering, demonstrated a heterogeneous pattern of VCAM-1 expression extending beyond the site of vessel injury. The morphology of the local inflammation patterns can have prognostic implications.[24] This high-resolution mapping of early stage inflammation would not have been possible with MRI alone as lesions are difficult to detect in small vessels against anatomical background. Whole body MRI screening in humans would be even more challenging. Thus, the combination of PET and MRI provided complementary functions in this imaging application.

Simultaneous PET/MRI imaging of the multimodal NP afforded a number of advantages over serial imaging including: 1) avoiding movement of the subject between scanner-, which allowed for facile spatial co-registration to do the lack of soft tissue movement experienced with scanning with separate instruments; 2) eliminating time delay between PET and MRI scans, which avoided complications with subject transport under anesthesia; 3) reduced exposure to personnel associated with handling and transporting the subject between instruments in different facilities. Given that many high-resolution MRI techniques are often time consuming and have a limited field of view, the availability of hybrid instrumentation can allow us to localize the ROI using PET without loss of spatial fidelity that would come with moving the subject to a separate scanner. This would be especially important in the clinical scenario whereby the site of injury may not be known a priori.

The approach of using hybrid PET/MRI instrumentation along with multimodal imaging probes provides a powerful platform for both research and clinical pursuits. From a research perspective, hybrid imaging can provide direct information about the efficacy and distribution of multimodal molecular imaging agents, providing vital information to refine and improve the signal characteristics of these agents. From a clinical perspective, PET/MRI hybrid instruments are available now in a limited number of locations and their optimal applications are under investigation. Oncological applications have been some of the earliest imaging studies to demonstrate beneficial results from hybrid PET/MRI.[39, 40] For example, a study of PET/MRI for clinical prostate cancer imaging found that PET/MRI may demonstrate improved sensitivity to metastatic lesions compared to PET/CT.[41] Hope for improved diagnoses through hybrid PET/MRI is high for neurological applications, where the temporal alignment of hybrid instruments could be an advantage for examining brain processes in real-time.[16]

Cardiovascular applications that could benefit from PET/MRI are still under investigation [22, 42]; however, the simultaneous acquisition capability could offer advantages for imaging fast moving hearts and arteries. Most reports with PET/MRI have focused primarily on unimodal contrast agents such as ^18^FDG using MRI as an anatomic reference.[15, 43, 44] We demonstrate that PET-guided MRI using multimodal agents can facilitate high-resolution visualization of molecular targets, and establish utility for mapping a marker associated with inflammation. This can serve as a powerful research tool for drug validation by allowing monitoring of biomarker targets. Early diagnosis and patient stratification by PET/MRI could provide clinical benefit as drugs to ameliorate plaque vulnerability come available.

In summary, we have demonstrated an approach to identify early inflammatory changes in vessel injury using a targeted multimodal probe coupled with hybrid PET/MRI imaging *in vivo*. Such approaches may be able to provide important insights into the pathophysiology and clinical management of vulnerable plaques. Furthermore, simultaneous PET/MRI allowed direct comparison of the PET and MRI signal, providing insights into the signal derived from the multimodal agent that can guide continued development of the instrument as well as the imaging agent.

## 5. Materials and Methods

### 5.1 Materials

Materials were obtained from commercial suppliers and used directly, unless otherwise noted. Dextran (from leuconostoc, average mol. wt. 9,000–11,000) and ferric chloride hexahydrate (FeCl_3_·6H_2_O, Fw 270.29 g/mol) were purchased from Sigma-Aldrich. Ferrous chloride tetrahydrate (FeCl_2_·4H_2_O, Fw 198.81 g/mol) was from Fluka. Ammonium hydroxide (28–30%), sodium bicarbonate, and sodium hydroxide were from Fisher Scientific. p-SCN-Bn-DOTA was from Macrocyclics, Inc (Dallas, TX). N-succinimidyliodoacetate (SIA), purchased from Pierce (Rockford, IL). Spectra/por^®^ dialysis membrane (mol. wt. cut-off 50,000) was acquired from Spectrum Laboratories, Inc. The VHPKQHR(MiniPEG1)C peptide was from Genscript (≥ 95% purity). Water was purified using a Millipore Milli-Q Synthesis purifier (18.0 MΩ cm, Barnstead).

### 5.2 Nanoparticle synthesis and characterization

#### 5.2.1 Synthesis of dextran coated iron oxide (DIO) nanoparticles

Dextran was reduced by published methods as summarized here and outlined in Figure 1.[25] Nanopure water (18.0 MΩ cm, Barnstead) was degassed with argon and used throughout the synthesis process. A mixture of dextran (molecular weight 10,000) and sodium borohydride (26 equivalents) was stirred at room temperature for 12 hours. The solution was adjusted to pH=7, dialyzed against degassed nanopure water and lyophilized to form reduced dextran as a white solid. A mixture of reduced dextran and FeCl_3_·6H_2_O in a molar ratio of 1:27 (total polysaccharide: FeCl_3_·6H_2_O) was dissolved in deionized nanopure water. The solution was bubbled with argon and cooled to 4 °C in an ice-water bath. Fe^2+^ solution was freshly prepared by dissolving FeCl_2_·4H_2_O in degassed water (with a Fe^3+^:Fe^2+^ ratio in a range from 1.47 to 1.5) and stored on ice. The Fe^2+^ solution was added to Fe^3+^ mixture using a syringe followed by adding chilled (4 °C) NH_4_OH (NH_4_OH: Fe^3+^ = 16:1) dropwise with vigorous stirring. The ice-water bath was removed and the mixture was heated to 85 °C and kept at 85 ± 5 °C for 2 hours (argon flow may stop 2–3 min after the temperature reaches 85 °C). After cooling to room temperature, the solution was dialyzed against deionized water in a dialysis bag with a molecular weight cut-off of 50,000 Da for 72 hours with 8-10 changes of water to remove reactants. The resulting product, DIO, was lyophilized and stored at 4 °C.[26]

#### 5.2.2 Cross-linking and amination of the DIO nanoparticles

The DIO nanoparticles were cross-linked and aminated for the attachment of two ligands: DOTA (chelator of radioactive ^64^Cu) and the VCAM-1 targeting peptide (Figure 1). The cross-linking and amination were performed as previously reported with slight modifications.[27] DIO (2.0g), NaOH (4.02g) pellets and deionized water (40mL) were added to a 100mL round-bottomed flask and stirred for 30 min. Then epichlorohydrin was added to the mixture and the solution was stirred for 24h. The solution was dialyzed against deionized water with 8-10 changes of deionized water and lyophilized to yield brown solid. The solid (2.0 g), together with ammonium hydroxide (250mL), was then transferred to a 500mL round-bottomed flask and stirred for 36h. Excess ammonium hydroxide was removed by dialyzing the solution against deionized water for 72h with 8-10 changes of deionized water.

#### 5.2.3 Conjugation of DOTA and VCAM-1 targeting peptide to the nanoparticle surface

DOTA was conjugated to the aminated DIO surface based on a literature method. Briefly, p-SCN-Bn-DOTA (6.71mg), aminated nanoparticle (135mg) and 0.1 M sodium borate buffer solution (2mL) were added to a 10 ml round-bottomed flask (Figure 1b)[28]. Approximately five drops of sodium hydroxide aqueous solution (1N) was used to bring the solution pH to 8.5. The pH of the mixture was monitored during the reaction. The mixture was stirred for 24h, then dialyzed against deionized water for 72h with 8-10 changes of water in a dialysis bag with a molecular weight cut-off of 50,000 Da and then lyophilized to give a brown solid of DOTA conjugated aminated DIO (DIO-DOTA). The conjugation of DOTA to the aminated DIO was confirmed by Fourier Transform Infrared (IR) Spectroscopy. DIO-DOTA was then used for anti-VCAM-1 peptide attachment based on previous report[29], described here briefly.

DIO-DOTA (22mg Fe in 1.5mL DMSO) was added to 0.5mL of 0.1M Na_2_HPO_4_ in water and 0.5mL of 15mM SIA in DMSO in a 10mL round-bottomed flask. The mixture was stirred for 1h at room temperature followed by another addition of 0.5mL of 15mM SIA (N-succinimidyliodoacetate) in DMSO. Iodoacetyl-DIO was separated from iodoacetic acid using a Sephadex G-25 column equilibrated with 0.025M citrate buffer pH 6.5 at 4 °C. Then 6-7 mg of polymer-modified peptide (VHPKQHR(MiniPEG1)C) in 0.6mL of citrate buffer was added to 5mL of iodoacetyl-DIO solution and the mixture was incubated overnight at room temperature. The purchased peptide, C_49_H_84_N_18_O_14_S_1_, had a molecular weight of 1181.37 Da and an isoelectric point of pH 9.84. MiniPEG1 (8-amino-3,6-dioxaoctanoic acid) spacer was inserted to distance the peptide from the surface of the nanoparticle to allow for proper tertiary folded structure and allow better VCAM binding. Unreacted peptide was removed by using a Sephadex G-25 column equilibrated with 0.025M citrate buffer pH 6.5 at 4°C. The purified solution was lyophilized to give the final product that is the DIO nanoparticle with two ligands (DOTA and VCAM-1 targeting peptide) on the surface (VDIO-DOTA).

#### 5.2.4 Copper-64 labeled VDIO-DOTA

Copper-64 was employed due to its relatively long half-life (12.7 h)[30] and comparatively stable coordination with multidentate macrocyclic compounds such as DOTA (log K_ML_ = 22.3).[31] VDIO-DOTA (25 mg) was dissolved in 150 µL of 0.2 M pH 5.5 sodium acetate-acetic acid buffer. Copper-64 (~ 2.5 mCi) was added to the vial and the mixture was vortexed for 5 seconds to obtain a uniform solution. The solution was incubated at 55-60 °C for 45 minutes. EDTA aqueous solution (16 µL, 100 mM) was added to the vial and vortexed for 5 seconds to get the solution uniform; then solution was incubated at 55-60 °C for 15 minutes. The crude product was purified by centrifuge filtration with 10K Da nanosep filtration tube (Millipore Inc., Billerica, MA, 30 min @14,000 rpm) and washed 3 times with pH 5.5 sodium acetate-acetic acid buffer. Each time the washing was removed by centrifuge filtration with a 10-kDa Nanosep filtration tube (10 min at 14,000 rpm). After three washings the filtration tube was turned over and inserted into a new vial, and the nanoparticles, VDIO-^64^Cu-DOTA, were collected by centrifuge (2 min @ 1,000 rpm). The radioactive nanoparticles were diluted to 650 µL with saline (0.9%) and radioactivity of the solution was measured with a Fluke Biomedical Dose calibrator (34-162 CAL/RAD MARK IV, Cleveland, OH). The radiolabeling yield (%) for the NP, percent incorporation of radioisotope, was determined by dividing the radioactivity of the collected NP by the total activity applied to the column and multiplying by 100. For injection to animals VDIO-^64^Cu-DOTA (~ 15 µL) was passed through a sterile 0.22-micron filter before use.

#### 5.2.5 Characterization of VDIO-DOTA nanoparticles

The iron concentration (mg) per unit mass of nanoparticles in DIO or VDIO-DOTA was measured with a Varian AA 220FS atomic absorption (AA) spectrophotometer using an air/acetylene flame. The iron-oxide core size of the VDIO-DOTA was measured by Transmission Electron Microscopy (TEM) on a Philips CM-12, operating at 80 kV and equipped with a GatanMegascan 795 digital camera. The core size was found by averaging the measurements of 500 particles after drying a dilute drop of VDIO-DOTA particles over a lamp on a copper grid. The average hydrodynamic particle size (mean volume diameter) and distribution was measured using Dynamic Light Scattering (DLS) with a Nanotrac 150 particle size analyzer (Microtrac, Inc., Montgomeryville, PA) and geometric eight-root regression, with no residuals, was used to fit the data. The nano-range option was selected and a scan time of 90 seconds was used.

Transverse relaxation times (*T*_2_) of the VDIO-DOTA particles were measured at 60 MHz (1.4 T) and 37 °C on a BrukerMinispec mq60. The relaxivity was given as the slope of the straight line plotted as the function of 1/*T*_2_ vs iron concentration. DIO and VDIO-DOTA were diluted in pH 7.0 deionized water to give five aqueous solutions (300 µL each): 10.5, 5.25, 2.625, 1.313, and 0.656 mg Fe/L, respectively. Iron concentrations in each dilution were determined using Atomic Absorption Spectroscopy. *T*_2_ values were measured using a Carr-Purcell-Meiboom-Gill (CPMG) sequence with τ = 1 ms, and 200 data points. Each solution was incubated at 37°C for 5 minutes before measurement.

### 5.3 Cytotoxicity

Cytotoxicity of VDIO-DOTA was evaluated on HepG2 liver cells using C_12_ – Resazurin viability assays. HepG2 liver cells were cultured and maintained in Minimum Essential Medium containing 10% FBS, 200 U/mL penicillin, 200 µg/mL streptomycin, 1 mM sodium pyruvate, and 1 mM nonessential amino acids at 37°C in a humidified 5% CO_2_ atmosphere. Cells were plated in 96-well dishes at a concentration of 1 x 10^4^ cells per well and incubated overnight (5% CO_2_, 37°C). After overnight incubation, media was replaced with fresh media containing VDIO-DOTA nanoparticles of varying concentration (0, 0.04, 0.2, 1, 4 and 10mM iron). Each concentration was performed in triplicate for statistical relevance. Cells were also treated with DIO nanoparticles under the same conditions as control. Cells were plated in three 96-well plates for measurements at time points of 4h, 24h and 48h. At each time point, nanoparticle solutions were removed and cells were washed with 1X PBS for three times. Then fresh media containing C_12_ – Resazurin (5µM) was added to each well of cells. After 15min of incubation, fluorescence was measured using a Safire monochromator microplate reader (Tecan Austria G.M.B.H., Austria) with excitation of 563nm and emission of 587nm.

### 5.4 Iron staining

Slides from the inflamed carotid were stained to evaluated for iron within the plaques using Prussian Blue Solution[32], a 1:1 mixture of 3% hydrochloride acid solution and 3% potassium ferrocyanide solution, to detect ferric ion (Fe^3+^) in the tissue. Any ferric ion present in the tissue reacted with the ferrocyanide and results in the formation of the bright blue pigment, Prussian blue, or ferric ferrocyanide. First, slides were deparaffinized by toluene and rehydrated through changes of ethanol with decreasing concentrations (100%, 95% and 75%). After rinsing, slides were placed in Prussian blue solution for 30 min followed by a rinsing step to remove excess staining. Then Nuclear Fast Red Staining was used to counterstain other tissue (pink) for 10 min. After dehydration with alcohol and clearing with toluene, tissue sections were placed under coverslips on slides with mounting media.

### 5.5 Immunohistochemistry (IHC)

The tissue sections were stained for inflammatory marker VCAM-1. The deparaffinization and dehydration process were the same as that of iron staining. Antigen retrieval was performed to unmask binding sites of the primary antibodies. Heat Induced Epitope Retrieval (HIER) was performed using a Decloaking Chamber. 500 ml of deionized water was added into the chamber. A plastic staining jar with tissue slides immersed in Antigen Retrieval Solution (Sigma 10X Tris-HCl buffer, pH10, product #T6455) was placed in the Decloaker chamber. The Decloaker was programed for 30 seconds at 125 °C followed by 10 seconds at 85 °C at 22.5 psi. After the heating process was done, the staining jar was removed from the Decloaker and cooled for 15 minutes followed by TBST rinse (Fisher 20X Tris buffered saline with Tween-20 used as 1X diluted solution). Then endogenous peroxidase block was performed by using 3% H_2_O_2_ in water to cancel the interference of peroxidase in the final step.[33] Protein block buffer (Dako Protein Block) was used to treat the tissue slides to mask the non-specific binding sites. Samples were labeled using an indirect method. Anti-VCAM-1 antibody (Rabbit monoclonal [EPR5047] to VCAM-1, Abcam Inc., MA) was used as primary antibody to detect the VCAM-1 expression. Ready-to-use polymers carrying horseradish peroxidase (Dako North America, Inc., Carpinteria, CA) were then used as the secondary antibody. In the final staining step, peroxidase on the polymers reacted with hydrogen peroxide to reduce the DAB (3,3‘-Diaminobenzidine) substrate and generate a brown product in regions of VCAM-1 expression. Slides were examined under microscope.

### 5.6 Animal Studies

#### 5.6.1 Animal model

All animal studies were performed under protocols approved by the Animal Care and Use Committee of the University of California, Davis and the California Institute of Technology. Female C57BL/6 ApoE−/− (10 weeks old, Jax West Laboratories, West Sacramento, CA) mice were used for the experiments as described in previous studies.[14] Eight days prior to imaging, the left carotid artery of each mouse was ligated. Eight days was chosen because prior studies showed that VCAM-1 expression peaks between 7 and 10 days post-ligation in ApoE−/− mice.[34] At this stage pronounced plaques have not yet formed, but inflammation is evident. To perform the ligation a medial incision was made between the mandible and clavicle, exposing the glands and vessels of the neck. The carotid artery was singled out from the surrounding tissue, while protecting and excluding the parallel-running vagus nerve. A 6/0 silk suture was threaded under the dorsal side of the carotid artery and was tied off to cause injury to the site. The procedure was concluded with five to six interrupted 4/0 Ethicon (Ethicon Inc) suture to re-approximate the skin of the original ventral incision. The mice were monitored twice a day for approximately four days to check for irritation and to administer analgesics when appropriate. Subsequent to ligation, mice were placed on a high fat diet for seven days. (TD 88137, Harlan Laboratories Inc, Madison, WI).

#### 5.6.2 *In vivo* MRI-only studies

MRI experiments were performed to determine the optimal injection dosages for the NPs prior to radiolabeling. Prior to NP injection, a pre-scan was taken as baseline. Then VDIO at dosages of 6 mg Fe/kg body mass, 30 mg Fe/kg and 60 mg Fe/kg, were injected intravenously via the tail vein catheter (N = 4 per concentration). Images were acquired at 4 and 24 hours post-injection.

All images were acquired on a 7T (Bruker Biospec) small animal scanner using a home built quadrature RF volume coil (Cleveland, OH). For all time points, the animal was anesthetized with a 1.5% isoflurane: air mixture and kept at 35–37 °C with warm air flowing through the bore while the respiration was monitored (MP150, Biopac, Goleta, CA). After localizing the region of interest (ROI) around the neck using a RARE spin echo sequence (TR/TE = 4000/22 ms, matrix size = 128 × 128, FOV = 35.35 × 35.35 mm^2^, slice thickness = 0.754 mm), the common carotid arteries were located with a time-of-flight angiography sequence with venous saturation (FL2D_ANGIO method, Paravision 4.0: TR/TE = 13.7/3.5 ms, matrix size = 150 × 100; zero-filled to 256 × 100, FOV = 30 × 20 mm^2^, slice thickness = 0.754 mm). A T_2*_ weighted multiple-gradient echo sequence was then utilized to visualize the uptake of nanoparticles (TR/TE = 718/3, 7, 11, 15, 19, 23 ms, F.A. = 25°, matrix size = 175 × 100; zero-filled to 234 × 133, FOV = 35 × 20 mm^2^, slice thickness = 0.754 mm) at the region of the common carotid arteries.

#### 5.6.3 *In vivo* PET/MRI imaging

Studies were performed using an integrated small animal PET/MRI system, consisting of a first-generation MR-compatible PET insert (constructed by Simon Cherry et. al at UC Davis) that is fitted within a 7T small animal MRI scanner. [16, 17] This enabled simultaneous PET/MRI images to be acquired.

Mice (N = 4) were surgically prepared as described above. Pre-scans were obtained with MRI as baseline. Next, mice were injected intravenously via the tail vein with 30 mg Fe/kg VDIO-^64^Cu-DOTA (~700uCi per mouse, 92% radiation yield) followed by a 150 µL of saline flush. The activity of the injected dose was confirmed by measuring the difference in radioactivity contained in the syringe before and after injection on a dose calibrator. Imaging was subsequently performed at 4 and 24 hours post-injection. MRI images were acquired identically to the MRI-only studies. PET images were acquired with scan duration of 600 seconds at the 4 hour time point and 300 seconds at the 24 hour time point. Images were reconstructed and co-registered to the MRI dataset as previously described.[17, 35]

To compare the focal NP uptake between time points, dosage and subjects, we calculated a contrast ratio (CR) metric as previously described.[36] This normalizes particle uptake at the ligation site between subjects and factors out the signal contribution due to blood borne particles by comparison with the contralateral control.

Briefly, CR is defined as:

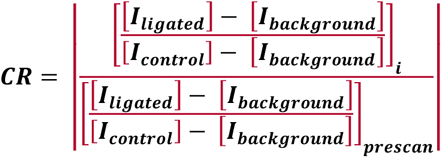

Where *I*_ligated_ is the mean intensity of the ROI drawn around the ligated carotid artery, *I*_control_ is the mean intensity of the ROI drawn around the contra-lateral carotid. *I*_background_ is the mean intensity of the ROI drawn in the spinal cord at the same image slice as the other ROIs and *i* is either 4 or 24 hours post-injection time points. ROIs were drawn manually at slice levels approximately located at the common carotid arteries. These were matched between time points. Angiography images were used to guide ROI delineation around the carotid arteries. Because previous reports noted that the carotid vessels along with the wall are ~1 mm in diameter,[36] all arterial ROIs had diameters of 1.5 mm.

Representative region of interests used for CR calculation are indicated in Figure 3. Values greater than 1 indicate localization at the ligation site. The T_2*_ weighted image sets at TE = 11 ms were used. All images were analyzed using ImageJ.

#### 5.6.4 Visualization of image volumes

We developed a hardware accelerated volume rendering system with enhanced rendering quality to provide better visualization of the PET and MRI signal at the site of the inflamed arteries. To provide anatomical context, we superimpose a cross section of the data. Illustrative rendering[37] is applied to this cross section for relative spatial position of the target vessel. The vessel ROIs were manually segmented from the MRI images to identify the vessel walls. A pre-integrated transfer function[38] was used to visualize the thin layer of vessel wall as a smooth and continuous surface. To highlight the NP uptake on the surface of the arteries, we used a color map to sort the surface MR signal ranging from red (low MR signal), through blue (medium) to light pink (high MR).

### 5.7 Histology

After the *in vivo* animal study, organs known to be involved in disease or clearance, i.e. hearts, carotid arteries, kidneys, spleens and livers, were collected and used for histology. The tissue was fixed in 4% formaldehyde, and then dehydrated by passing tissue through increasing concentrations of ethanol (75%, 95% and 100%). Then the tissue was placed in warm paraffin wax, and the melted wax filled the spaces that used to contain water. After cooling, the tissue hardened into a paraffin block from which 5-micron tissue slices were sectioned and mounted on glass slides. The tissue sections were stained for iron and VCAM-1 as detailed in Supplemental Information.

## Supplementary Materials

N/A

### Acknowledgments

The authors wish to thank F. Hayes and P. Kysar for their help with TEM and P. Hrvatin for help with AA spectroscopy. We thank Andre Jefferson and Haick Issian of the Caltech Radiation Safety office for help with the radiation studies and Naomi Santa-Maria for technical assistance in imaging. We thank Simon Cherry and his group for technical assistance with the hybrid imaging insert. The authors wish to acknowledge the National Institutes of Health (EB008576-01 and EB000993), the Center for Molecular and Genomic Imaging at the University of California, Davis (U24 CA 110804), and the NMR award of the University of California, Davis for support of this work.

### Author Contributions

AYL conceived and designed experiments; SD, TN, AH, TT, CT contributed to experiment design, performed experiments and contributed to data analysis; LZ prepared computer programs to render the imaging data; JP, IE performed and provided training for histology; REJ, KP contributed to experimental design and data analysis.

## References

1. Fuster V, Lois F, Franco M. Early identification of atherosclerotic disease by noninvasive imaging. Nature Reviews Cardiology. 2010;7(6):327–33.

2. Finn AV, Nakano M, Narula J, Kolodgie FD, Virmani R. Concept of vulnerable/unstable plaque. Arteriosclerosis, thrombosis, and vascular biology. 2010;30(7):1282–92.

3. Kolodgie FD, Burke AP, Skorija KS, Ladich E, Kutys R, Makuria AT, et al. Lipoprotein-associated phospholipase A2 protein expression in the natural progression of human coronary atherosclerosis. Arteriosclerosis, thrombosis, and vascular biology. 2006;26(11):2523–9.

4. Braganza D, Bennett M. New insights into atherosclerotic plaque rupture. Postgraduate medical journal. 2001;77(904):94–8.

5. DeMarco JK, Huston III J. Imaging of high-risk carotid artery plaques: current status and future directions. Neurosurgical focus. 2014;36(1):E1.

6. Cai J-M, Hatsukami TS, Ferguson MS, Small R, Polissar NL, Yuan C. Classification of human carotid atherosclerotic lesions with in vivo multicontrast magnetic resonance imaging. Circulation. 2002;106(11):1368–73.

7. Hyafil F, Schindler A, Sepp D, Obenhuber T, Bayer-Karpinska A, Boeckh-Behrens T, et al. High-risk plaque features can be detected in non-stenotic carotid plaques of patients with ischaemic stroke classified as cryptogenic using combined F-18-FDG PET/MR imaging. European Journal of Nuclear Medicine and Molecular Imaging. 2016 Feb;43(2):270–9.

8. Moon SH, Cho YS, Noh TS, Choi JY, Kim BT, Lee KH. Carotid FDG Uptake Improves Prediction of Future Cardiovascular Events in Asymptomatic Individuals. Jacc-Cardiovascular Imaging. 2015 Aug;8(8):949–56.

9. van der Valk FM, Verweij SL, Zwinderman KAH, Strang AC, Kaiser Y, Marquering HA, et al. Thresholds for Arterial Wall Inflammation Quantified by F-18-FDG PET Imaging. Jacc-Cardiovascular Imaging. 2016 Oct;9(10):1198–207.

10. Yarasheski KE, Laciny E, Overton ET, Reeds DN, Harrod M, Baldwin S, et al. (18)FDG PET-CT imaging detects arterial inflammation and early atherosclerosis in HIV-infected adults with cardiovascular disease risk factors. J Inflamm (Lond). 2012;9:26.

11. Huet P, Burg S, Le Guludec D, Hyafil F, Buvat I. Variability and Uncertainty of F-18-FDG PET Imaging Protocols for Assessing Inflammation in Atherosclerosis: Suggestions for Improvement. Journal of Nuclear Medicine. 2015 Apr;56(4):552–9.

12. van der Valk FM, Verweij SL, Zwinderman KA, Strang AC, Kaiser Y, Marquering HA, et al. Thresholds for Arterial Wall Inflammation Quantified by 18F-FDG PET Imaging: Implications for Vascular Interventional Studies. JACC Cardiovasc Imaging. 2016 Oct;9(10):1198–207.

13. Brammen L, Steiner S, Berent R, Sinzinger H. Molecular imaging of atherosclerotic lesions by positron emission tomography - can it meet the expectations? Vasa-European Journal of Vascular Medicine. 2016;45(2):125–32.

14. Jarrett BR, Correa C, Ma KL, Louie AY. In vivo mapping of vascular inflammation using multimodal imaging. PloS one. 2010;5(10):e13254.

15. Schindler TH. Cardiovascular PET/MR imaging: Quo Vadis? J Nucl Cardiol. 2016 Sep 22.

16. Catana C, Procissi D, Wu Y, Judenhofer MS, Qi J, Pichler BJ, et al. Simultaneous in vivo positron emission tomography and magnetic resonance imaging. Proceedings of the National Academy of Sciences. 2008;105(10):3705–10.

17. Ng TS, Bading JR, Park R, Sohi H, Procissi D, Colcher D, et al. Quantitative, simultaneous PET/MRI for intratumoral imaging with an MRI-compatible PET scanner. Journal of Nuclear Medicine. 2012;53(7):1102–9.

18. Nahrendorf M, Jaffer FA, Kelly KA, Sosnovik DE, Aikawa E, Libby P, et al. Noninvasive vascular cell adhesion molecule-1 imaging identifies inflammatory activation of cells in atherosclerosis. Circulation. 2006;114(14):1504–11.

19. Broisat A, Toczek J, Dumas LS, Ahmadi M, Bacot S, Perret P, et al. Tc-99m-cAbVCAM1-5 Imaging Is a Sensitive and Reproducible Tool for the Detection of Inflamed Atherosclerotic Lesions in Mice. Journal of Nuclear Medicine. 2014 Oct;55(10):1678–84.

20. Dimastromatteo J, Broisat A, Perret P, Ahmadi M, Boturyn D, Dumy P, et al. In Vivo Molecular Imaging of Atherosclerotic Lesions in ApoE(-/-) Mice Using VCAM-1-Specific, Tc-99m-Labeled Peptidic Sequences. Journal of Nuclear Medicine. 2013 Aug;54(8):1442–9.

21. Wu J, Leong-Poi H, Bin J, Yang L, Liao Y, Liu Y, et al. Efficacy of Contrast-enhanced US and Magnetic Microbubbles Targeted to Vascular Cell Adhesion Molecule–1 for Molecular Imaging of Atherosclerosis. Radiology. 2011;260(2):463–71.

22. Ripa RS, Kjær A. Imaging Atherosclerosis with Hybrid Positron Emission Tomography/Magnetic Resonance Imaging. BioMed Research International. 2015;2015:8.

23. O’Brien J, Wilson I, Orton T, Pognan F. Investigation of the Alamar Blue (resazurin) fluorescent dye for the assessment of mammalian cell cytotoxicity. European Journal of Biochemistry. 2000;267(17):5421–6.

24. Petersen C, Peçanha PB, Venneri L, Pasanisi E, Pratali L, Picano E. The impact of carotid plaque presence and morphology on mortality outcome in cardiological patients. Cardiovasc Ultrasound. 2006;4(16).

25. Wilson CM. Synthesis of Ultrasmall Superparamagnetic Iron Oxide Nanoparticles for fMRI.

26. Jarrett BR, Frendo M, Vogan J, Louie AY. Size-controlled synthesis of dextran sulfate coated iron oxide nanoparticles for magnetic resonance imaging. Nanotechnology. 2007;18(3):035603.

27. Wunderbaldinger P, Josephson L, Bremer C, Moore A, Weissleder R. Detection of lymph node metastases by contrast-enhanced MRI in an experimental model. Magnetic resonance in medicine. 2002;47(2):292–7.

28. Tu C, Ng TS, Jacobs RE, Louie AY. Multimodality PET/MRI agents targeted to activated macrophages. JBIC Journal of Biological Inorganic Chemistry. 2014;19(2):247–58.

29. Josephson L, Kircher MF, Mahmood U, Tang Y, Weissleder R. Near-infrared fluorescent nanoparticles as combined MR/optical imaging probes. Bioconjugate chemistry. 2002;13(3):554–60.

30. Gustafsson B, Youens S, Louie AY. Development of contrast agents targeted to macrophage scavenger receptors for MRI of vascular inflammation. Bioconjugate chemistry. 2006;17(2):538–47.

31. Anderegg G, Arnaud-Neu F, Delgado R, Felcman J, Popov K. Critical evaluation of stability constants of metal complexes of complexones for biomedical and environmental applications+(IUPAC Technical Report). Pure and applied chemistry. 2005;77(8):1445–95.

32. Sheehan DC, Hrapchak BB. Theory and practice of histotechnology: Mosby St. Louis; 1980.

33. Streefkerk J. Inhibition of erythrocyte pseudoperoxidase activity by treatment with hydrogen peroxide following methanol. Journal of Histochemistry & Cytochemistry. 1972;20(10):829–31.

34. Barringhaus KG, Phillips JW, Thatte JS, Sanders JM, Czarnik AC, Bennett DK, et al. α4β1 Integrin (VLA-4) Blockade Attenuates both Early and Late Leukocyte Recruitment and Neointimal Growth following Carotid Injury in Apolipoprotein E (–/–) Mice. Journal of vascular research. 2004;41(3):252–60.

35. Ng TS, Procissi D, Wu Y, Jacobs RE. A robust coregistration method for in vivo studies using a first generation simultaneous PET/MR scanner. Medical physics. 2010;37(5):1995–2003.

36. Tu C, Ng TS, Sohi HK, Palko HA, House A, Jacobs RE, et al. Receptor-targeted iron oxide nanoparticles for molecular MR imaging of inflamed atherosclerotic plaques. Biomaterials. 2011;32(29):7209–16.

37. Lum EB, Ma K-L, editors. Hardware-accelerated parallel non-photorealistic volume rendering. Proceedings of the 2nd international symposium on Non-photorealistic animation and rendering; 2002: ACM.

38. Lum EB, Wilson B, Ma K-L, editors. High-quality lighting and efficient pre-integration for volume rendering. Proceedings of the Sixth Joint Eurographics-IEEE TCVG conference on Visualization; 2004: Eurographics Association.

39. Bashir U, Mallia A, Stirling J, Joemon J, MacKewn J, Charles-Edwards G, et al. PET/MRI in Oncological Imaging: State of the Art. Diagnostics. 2015;5(3):333–57.

40. Partovi S, Kohan A, Rubbert C, Vercher-Conejero JL, Gaeta C, Yuh R, et al. Clinical oncologic applications of PET/MRI: a new horizon. Am J Nucl Med Mol Imaging. 2014;4(2):202–12.

41. Catalano OA, Rosen BR, Sahani DV, Hahn PF, Guimaraes AR, Vangel MG, et al. Clinical impact of PET/MR imaging in patients with cancer undergoing same-day PET/CT: initial experience in 134 patients--a hypothesis-generating exploratory study. Radiology. 2013 Dec;269(3):857–69.

42. Nensa F, Schlosser T. Cardiovascular hybrid imaging using PET/MRI. Rofo. 2014 Dec;186(12):1094–101.

43. Li X, Heber D, Rausch I, Beitzke D, Mayerhoefer ME, Rasul S, et al. Quantitative assessment of atherosclerotic plaques on 18F-FDG PET/MRI: comparison with a PET/CT hybrid system. Eur J Nucl Med Mol Imaging. 2016;43:1503–12.

44. Masuda A, Yamaki T, Sakamoto N, Kunii H, Ito H, Nanbu T, et al. Vulnerable plaque on the common iliac artery detected by F-18-FDG PET/MRI. European Journal of Nuclear Medicine and Molecular Imaging. 2016 Apr;43(4):793–4.

